# DEK::NUP214 acts as an XPO1-dependent transcriptional activator of essential leukemia genes

**DOI:** 10.1101/2025.03.05.641116

**Authors:** Fadimana Kaya, Findlay Bewicke-Copley, Juho J Miettinen, Pedro Casado, Eve Leddy, Özgen Deniz, Vincent-Philippe Lavallée, Celine Philippe, Jiexin Zheng, Florian Grebien, Naeem Khan, Szilvia Krizsán, Joseph Saad, Alexis Nolin-Lapalme, Josée Hébert, Sébastien Lemieux, Eric Audemard, Janet Matthews, Marianne Grantham, Doriana Di Bella, Krister Wennerberg, Alun Parsons, John Gribben, James D. Cavenagh, Sylvie D Freeman, Csaba Bödör, Guy Sauvageau, Jun Wang, Pilar Llamas-Sillero, Jean-Baptiste Cazier, David C Taussig, Dominique Bonnet, Pedro R Cutillas, Caroline A Heckman, Jude Fitzgibbon, Kevin Rouault-Pierre, Ana Rio-Machin

**Author notes:** **Correspondence:** /.

## Abstract

The t(6;9)(p22.3;q34.1) translocation/DEK::NUP214 fusion protein defines a distinct subgroup of younger AML patients classified as a separate disease entity by the World Health Organization. DEK is a nuclear factor with multifunctional roles, including gene regulation, while its fusion partner, NUP214, plays a pivotal role in nuclear export by interacting with transport receptors such as XPO1. However, the precise mechanism by which DEK::NUP214 drives leukemia remains unclear. A comprehensive multi-omics comparison of 57 AML primary samples (including whole genome sequencing, targeted sequencing, transcriptomics, and drug screening with > 500 compounds) revealed that t(6;9) cases display a selective response to XPO1 inhibitors (Selinexor & Eltanexor) and a distinct transcriptomic signature characterized by the overexpression of *FOXC1* and *HOX* genes that are key leukemia mediators. CUT&RUN experiments demonstrated the direct binding of DEK::NUP214 to the promoters of *FOXC1* and *HOXA/B* clusters. Strikingly, the expression of these genes and the binding of DEK::NUP214 to their regulatory regions were selectively reduced upon XPO1 inhibition in t(6;9) cells. Altogether, these results identified a novel function of DEK::NUP214 as an XPO1-dependent transcriptional activator of key leukemia drivers and provide a rationale to explore the use of XPO1 inhibitors in this patient population.

## Introduction

Acute myeloid leukemia (AML) with t(6;9)/DEK::NUP214 is recognized as a separate entity in the World Health Organization classification of myeloid neoplasms, accounting for 1% of all AML cases (1) and characterized by a high relapse rate and young age at diagnosis (median age of 23) (2). Co-occurrence with *FLT3*-ITD is observed in 70% of cases, which confers inferior outcomes with a 5-year overall survival rate <25%. The t(6;9)(p22.3;q34.1) chromosomal rearrangement results in the *in-frame* fusion of almost the entire peptide sequence of *DEK* with the C-terminal region of *NUP214*, encoding the chimeric protein DEK::NUP214 (2, 3). NUP214 is an FG nucleoporin anchored to the cytoplasmic ring of the Nuclear Pore Complex that interacts with multiple transport receptors, including exportin 1 (XPO1), and these interactions are maintained in DEK::NUP214 (3, 4). DEK is a ubiquitously expressed nuclear factor with multiple functions, including gene regulation (5). Overexpression of DEK::NUP214 induces proliferation of myeloid cells (6) and leads to the development of leukemia in mice (7). A recent study has shown that t(6;9) AML displays a gene expression signature reminiscent of other AML subtypes, such as *NPM1*-mutated patients, which is characterized by the overexpression of *HOX* genes, a key leukemia driver event (8). However, the molecular mechanisms underpinning the pathogenicity of DEK::NUP214 and how these transcriptional changes are orchestrated remain unclear. Here, through multi-omics and functional approaches in primary AML patient samples and human cellular models, we observed a strong and selective sensitivity to XPO1 inhibition of t(6;9) cells and demonstrated that DEK::NUP214 acts as an XPO1-dependent transcriptional activator of essential leukemia drivers.

### Study design

As part of a cohort of 57 cytogenetically poor-risk AML patients, four t(6;9) AML patient samples were subjected to a multi-omics approach, including RNA-sequencing, whole genome/targeted sequencing, and *in vitro* drug screening (> 500 compounds). RT-qPCR/RNA-seq and Western blotting assessed RNA and protein expression, respectively. Lentiviral shRNAs were used to target *FOXC1* in the FKH-1 (t(6;9) AML cell line), followed by functional assessments, including cell cycle, apoptosis, and colony-forming unit assays. CUT&RUN experiments revealed DEK::NUP214’s genomic binding sites. Detailed methodology is available in the supplemental data.

## Results and Discussion

### AML primary samples with t(6;9)/DEK::NUP214 display a unique transcriptional signature

In order to identify potential mediators underlying DEK::NUP214 leukemogenesis, we analyzed RNA-seq data of a cohort of 57 cytogenetically poor-risk AML samples that included four t(6;9) patients (supplemental Table 1) (9, 10). The median age of these DEK::NUP214 cases was 43.5 years (29–55 years) with a mean overall survival of 1.18 years (0.90–1.85 years). The mutation profile of each patient is shown in supplemental Table 2 with all 4 patients harboring a *FLT3*-ITD.

In an unsupervised RNA-seq analysis, we first observed that AML samples from t(6;9) patients clustered distinctly from other AMLs (Figure 1A), suggesting a unique transcriptional signature. To explore this further, we integrated our RNA-seq data with a separate series of 691 AML cases from all cytogenetic groups that included three additional t(6;9) cases (Leucegene project, https://leucegene.ca/). Our analysis identified 166 differentially expressed genes (p < 0.05, Fold Change > 1.5) in t(6;9) patients compared to other AML samples (Figure 1B; supplemental Table 3). Among the 104 significantly upregulated transcripts, we found known leukemia drivers (*FOXC1* and *HOXB* genes), genes previously reported as being dysregulated in t(6;9) patients (*EYA3, SESN1, NFIX* and *PRDM2*) (8, 11, 12) and novel genes such as *GGT5* (Figure 1B).

**Figure 1.**
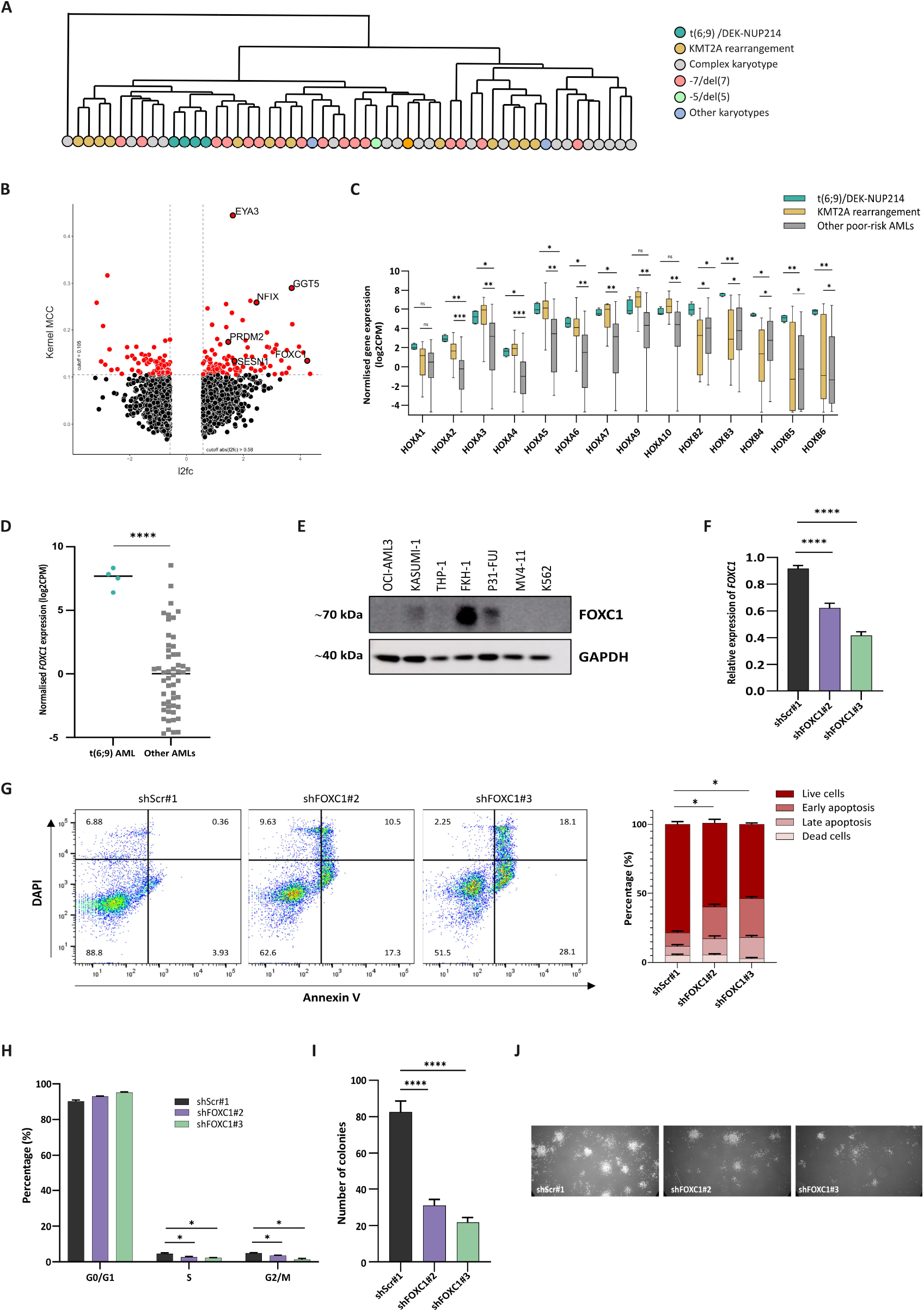
AML primary samples with t(6;9)/DEK::NUP214 display a unique transcriptional signature. **(A)** RNA-seq unsupervised clustering of 57 cytogenetically poor-risk AML samples (10), including four t(6;9) cases. Different colors represent distinct cytogenetic AML subtypes. **(B)** Volcano plot representing differentially expressed genes in t(6;9) primary AML samples vs other cytogenetic AML subtypes (RNA-seq analysis integrating our cohort of 57 poor-risk cases and 691 cases from the Leucegene cohort (Leucegene project, https://leucegene.ca/) (significance determined by Kernel MCC > 0.105 and log_2_FoldChange (l2fc) > 0.58). *Kernel MCC* is a shorthand for *Matthews Correlation Coefficient of a kernel density estimation (KDE)-based classifier*. **(C)** mRNA expression levels *HOXA* and *HOXB* genes in t(6;9) samples compared to *KMT2A*-rearranged and other poor-risk AMLs (n=57). **(D)** mRNA expression level of *FOXC1* in t(6;9) patients compared to other poor-risk AMLs (n=57). **(E)** FOXC1 protein expression across the AML cell line panel (OCI-AML3, KASUMI-1, THP-1, FKH-1, P31-FUJ, MV4;11 and K562). GAPDH was used as an endogenous control. **(F)** Bar chart showing *FOXC1* silencing efficiency by RT-qPCR after shRNA/scramble transduction of FKH1 cells. Relative expression levels were calculated using the ΔCT method, normalized to the average of GAPDH and 18S rRNA. **(G)** Representative flow cytometry plot for the apoptosis assay, with *FOXC1* knockdown (KD)/scramble (scr) FKH-1 cells stained with Annexin V (X-axis) and DAPI (Y-axis). The bar graph illustrates the percentage of live cells (DAPI^-^ Annexin V^-^), early apoptosis (DAPI^-^ Annexin V^+^), late apoptosis (DAPI^+^ Annexin V^+^), and dead cells (DAPI^+^ Annexin V^-^). **(H)** Bar graph representing cell cycle analysis by flow cytometry based on DNA content stained with DAPI of *FOXC1* KD/scr FKH-1 cells. Each group of bars indicates the percentage distribution of cells in G0/G1, S, and G2/M phases. **(I)** Bar graph showing the number of colonies of *FOXC1* KD/scr FKH-1 cells after the colony forming unit (CFU) assay. **(J)** Representative images of colonies corresponding to the CFU assay. Asterisks indicate statistical significance (*: p<0.05, **: p<0.01, ***: p<0.001, ****: p<0.0001 and ‘ns’ denotes non-significant differences).

Taking advantage of the FKH-1 AML cell line that harbors an identical t(6;9) translocation as the patients in our cohort (supplemental Figure 1A), we validated the significant up-regulation of the fore-mentioned genes in FKH-1 cells compared to a panel of 6 non-t(6;9) AML cell lines (supplemental Figure 1B).

*HOXA* and *HOXB* genes are a family of hematopoietic transcription factors whose deregulation is a feature of leukemogenesis. t(6;9) AML displays the overexpression of *HOXA* genes characteristic of other AML subtypes, such as patients with *NPM1* mutations or *KMT2A* rearrangements (Figure 1C) (8), whereas overexpression of *HOXB* genes appears characteristic of t(6;9) samples (Figure 1C). *FOXC1*, another transcription factor whose overexpression confers a monocyte/macrophage lineage differentiation block and is associated with AML poorer outcomes (13), is strikingly overexpressed (Log2FC = 4.2465) in t(6;9) cases compared to other AML samples (Figure 1D). This was also confirmed in the t(6;9) FKH-1 cell line at the protein level where FOXC1 was strongly expressed, while it was barely detected in other AML cells (Figure 1E). Given the established role of *FOXC1* in AML pathogenesis, we focused our attention on this transcription factor, noting that its shRNA-mediated silencing in FKH-1 cells (Figure 1F) led to a significant increase in apoptosis (Figure 1G), cell cycle arrest (Figure 1H), and a decrease in the number of colonies under hypoxic conditions (3% O_2_) (Figure 1I and 1J), paralleled to a significant downregulation of *HOX* genes expression (supplemental Figure 1C). These findings demonstrate an important role for *FOXC1* in the maintenance of t(6;9) leukemic cells, making it an attractive therapeutic target for the treatment of this disease subtype.

### XPO1 inhibition specifically rewires DEK::NUP214 transcriptional signature

In order to identify therapeutic vulnerabilities of t(6;9) patients, our four primary samples were subjected to an *in vitro* drug screening using our established drug sensitivity and resistance-testing platform (DSRT) (10). The complete drug panel comprises 527 licensed/investigational compounds, but smaller panels were assessed in patients P2 and P3, where the number of viable cells available was limiting. DSRT with the same drug panel had previously been applied to a cohort of 145 AML primary samples spanning all cytogenetic groups (10, 14) (supplemental Table 4). We integrated these existing data with our results in t(6;9) samples to identify the most selective and effective compounds for these patients. This analysis revealed that the two XPO1 inhibitors included in the drug screening, Eltanexor and Selinexor, ranked in the top-4 compounds tested (Figure 2A; supplemental Table 5). Likewise, FKH-1 showed enhanced sensitivity to XPO1 inhibitors compared to AML cell lines with distinct cytogenetic backgrounds (supplemental Figure 2A). In support of these findings, previous studies reported the interaction between DEK::NUP214 and XPO1 through the NUP214-FG domain (11) and, of note, the sole t(6;9) patient included in a phase I study of Selinexor in AML achieved complete remission (15). In order to investigate whether XPO1 inhibition alters t(6;9)/DEK::NUP214 transcriptomic profile, we performed RNA-seq post *in vitro* exposure to Selinexor of t(6;9) and non-t(6;9) primary AML cells. We included a treatment control, Omacetaxine, another compound with increased toxicity in t(6;9) cells identified during the drug screening (Figure 2A; supplemental Figure 2B). 36 out of the 104 genes upregulated at diagnosis in t(6;9) primary samples as compared to other AML samples (supplemental Table 3), including *FOXC1, NFIX, EYA3, SESN1, PRDM2* and *HOX* genes, were significantly downregulated in t(6;9) cells upon XPO1 inhibition (Figure 2B; supplemental Figure 2C; supplemental Table 6), but, critically, not in non-t(6;9) primary samples (supplemental Figure 2D; supplemental Table 6). No differences were observed in the expression of these genes upon Omacetaxine treatment in any of the samples analyzed (supplemental Figures 3A-C; supplemental Tables 6-7). These results suggest a strong XPO1 dependency in t(6;9) AML to establish its onco-transcriptomic signature.

**Figure 2.**
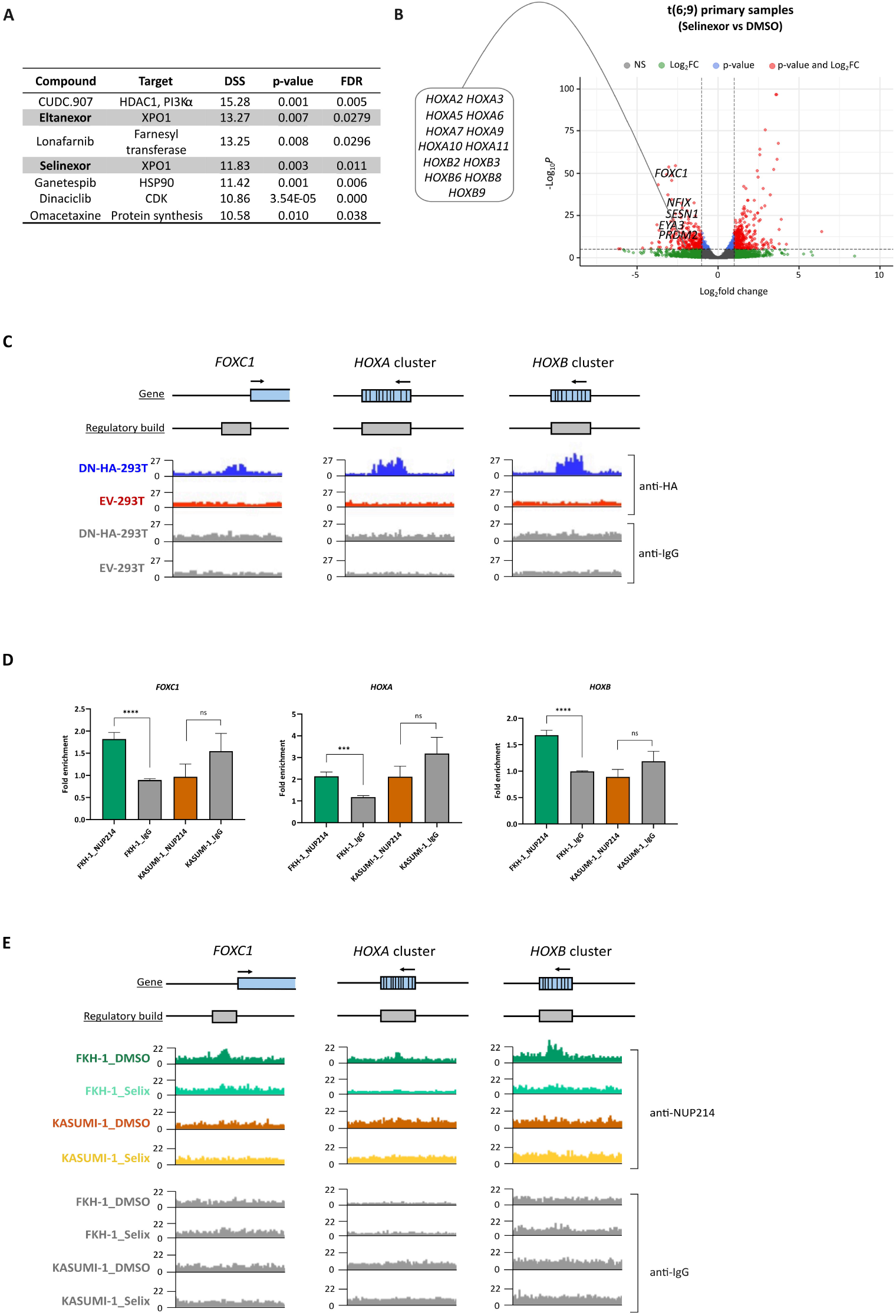
DEK::NUP214 specifically binds to the regulatory regions of key leukemia genes in an XPO1-dependent manner. **(A)** *In vitro* drug screening using a 527-drug panel in 4 t(6;9) patients and 145 primary AML samples from other cytogenetic groups (10, 14). The table shows the top-ranked compounds most specific and efficient for t(6;9) patients, with the XPO1 inhibitors (Selinexor and Eltanexor) highlighted in grey. **(B)** Volcano plot representing differentially expressed genes in two t(6;9) primary AML samples treated with Selinexor vs DMSO treatment control (RNA-seq results). Genes highlighted in the plot were significantly downregulated (p<0.05) after XPO1 inhibition. **(C)** IGV genome browser plots representing the binding peaks on the *FOXC1, HOXA* and *HOXB* promoters corresponding to the CUT&RUN experiments performed in DN-HA-293T and EV-293T cells using anti-HA antibody (and anti-IgG as a control). **(D)** CUT&Run-qPCR results show fold enrichment at the promoter regions of the *FOXC1, HOXA* and *HOXB* genes clusters in the t(6;9) AML FKH-1 and non-t(6;9) AML KASUMI-1 cells, compared to their respective IgG controls. FKH-1_NUP214 and KASUMI-1_NUP214 refer to CUT&Run performed with the NUP214 antibody, while FKH-1_IgG and KASUMI-1_IgG indicate the corresponding controls performed with the IgG antibody. Fold enrichment was calculated relative to the FKH-1_IgG sample. Statistical significance is indicated as ***: p<0.001, ***: p<0.0001, and ns: non-significant differences. (n=3). **(E)** IGV genome browser plots representing the binding peaks on the *FOXC1, HOXA* and *HOXB* promoters corresponding to the CUT&Run experiments performed in FKH-1 and Kasumi-1 cells after treatment with 200nM Selinexor for 48 hours and using anti-NUP214 antibody (and anti-IgG as a control) (*Selix:* treated with Selinexor; *DMSO*: treated with DMSO).

### DEK::NUP214 acts as an XPO1-dependent transcription factor

Our findings raised the question of whether DEK::NUP214 has a direct role in transcriptional regulation and whether XPO1 contributes to this effect. To address this, we first overexpressed HA tagged-DEK::NUP214 by lentiviral transduction of HEK-293T (DN-HA-293T model) and validated the interaction between DEK::NUP214 and XPO1 by co-immunoprecipitation (supplemental Figures 4A-B). We next conducted CUT&Run experiments in both the DN-HA-293T model (HA antibody) and the FKH-1 cell line (NUP214 antibody) using empty vector-HEK-293T model (EV-293T) and KASUMI-1 cell line as negative controls, respectively. These experiments revealed that DEK::NUP214 binds to the promoters of 801 genes (supplemental Table 8), with around one third encoding DNA or RNA binding proteins (FDR = 6.53e-09, GO:0003676) involved in transcription, DNA replication, DNA repair and RNA processing. Of note, 10 of the genes differentially expressed between t(6;9) and non-t(6;9) patients were also among the direct targets of DEK::NUP214 identified in the CUT&Run experiments, including *FOXC1* and *HOXB* genes (Figure 2C; supplemental Figure 4C). We also demonstrated that DEK::NUP214 binds to the promoters of the *HOXA* cluster (Figure 2C; supplemental Figure 4C). The binding of DEK-NUP214 to the regulatory regions of *FOXC1, HOXB* and *HOXA* genes was further confirmed by CUT&RUN-qPCR (16) (see Supplemental Material & Methods) in the FKH-1 cell line, using KASUMI-1 cells as a negative control (Figure 2D). These findings provide the first evidence of DEK::NUP214’s direct interaction with DNA, suggesting a role as a putative transcriptional activator of key leukemia players. CUT&Run experiments after Selinexor treatment of FKH-1 cells showed that XPO1 inhibition resulted in the loss of the binding of DEK::NUP214 to the regulatory regions of these target genes (Figure 2E), indicating that this interaction is dependent on the presence of XPO1 and explaining the specific gene expression rewiring after Selinexor treatment in t(6;9) cells.

Altogether, we demonstrate that a distinct transcriptional program is conserved in t(6;9) AML and characterized by the overexpression of essential leukemic factors (e.g., *FOXC1, HOXA* and *HOXB* genes) that are directly regulated by DEK::NUP214. We also found a functional interplay between XPO1 and the fusion protein, resulting in a selective sensitivity of t(6;9) cells to XPO1 inhibition that specifically reverts this transcriptomic profile.

## Contributions

AR-M and JF conceived the project. AR-M, JF, FK and KR-P designed the experiments and wrote the original draft. FK and AR-M performed research and collected data. FB-C, AR-M, FK, EL, OZ, VPL, NK, SK, AN-L, JH, SL, EA, PC and CB performed data analysis. JM, JS, KW, AP and CH designed, performed and analyzed the drug sensitivity screening. CP, JZ, NK, SK and DDB contributed to the experiments. JM and MG contributed to data collection. FG, JG, JC, SF, CG, GS, JW, JBC, DT, DB, PRC and CAH contributed to the conceptualization and with resources. Funding acquisition: AR-M, JF and FK. All authors reviewed the manuscript.

## Acknowledgements

The authors thank the patients for donating specimens for research to this study, their hematologists for the contribution and the tissue bank of the Barts Cancer Institute for collection and storage. This study was predominantly funded by the European Hematology Association through the AR-M’s EHA Junior Research Grant (EHA RG71), and Cancer Research UK (C15966/A24375). FK was supported by the Republic of Turkey Ministry of National Education, General Directorate of Higher and Foreign Education, Turkish Government Study Abroad Program. CP and KR-P were supported by the Barts Charity (G-002167) and the Kay Kendall Leukaemia fund (KKL1149). CB was funded by the National Research, Development, and Innovation Office, Hungary (grants K21-137948, TKP2021-EGA-24 and TKP2021-NVA-15) and the Momentum Program of the Hungarian Academy of Sciences. RNA sequencing of the Leucegene cohort was supported by Genome-Canada and Genome-Quebec. Leukemic samples and cytogenetic data of Leucegene cohort were provided by the Quebec leukemia cell bank supported by grants from the Cancer Research Network of the Fonds de recherche du Québec–Santé.

## Notes

**Competing Interests statement:** Authors have no conflicts of interest to disclose.

https://www.ncbi.nlm.nih.gov/geo/query/acc.cgi?acc=GSE279056

### Competing Interest Statement

The authors have declared no competing interest.

