## Supplemental figures, tables and methods for "DEK::NUP214 acts as an XPO1-dependent transcriptional activator of essential leukemia genes"

1 Centre for Haemato-Oncology, Barts Cancer Institute, Queen Mary University of London, United Kingdom; 2 Institute for Molecular Medicine Finland – FIMM, HiLIFE – Helsinki Institute of Life Science, iCAN Digital Precision Cancer Medicine Flagship, University of Helsinki, Helsinki, Finland; 3 Centre for Centre for Cancer Evolution, Barts Cancer Institute, Queen Mary University of London, London, United Kingdom; 4 Centre for Epigenetics, Queen Mary University of London, London, E1 2AT, United Kingdom; 5 Division of Hematology and Oncology, Centre Hospitalier Universitaire Sainte-Justine, Montréal, Canada; 6 INSERM U1242, University of Rennes, Rennes, France; Centre de Lutte contre le cancer Eugène Marquis, Rennes, France; 7 University College London Hospitals NHS Foundation Trust, London, United Kingdom; 8 Department of Biological Sciences and Pathobiology, University of Veterinary Medicine, Vienna, Austria; St. Anna Children's Cancer Research Institute (CCRI), Vienna, Austria; CeMM Research Center for Molecular Medicine of the Austrian Academy of Sciences, Vienna, Austria; 9 School of Infection, Inflammation and Immunology, University of Birmingham College of Medicine and Health, United Kingdom; 10 HCEMM-SU, MTA-SE „Lendület" Molecular Oncohematology Research Group, Department of Pathology and Experimental Cancer Research, Semmelweis University, Budapest, Hungary; 11 Institute for Research in Immunology and Cancer, Montreal, Canada; 12 Institut universitaire d'hémato-oncologie et de thérapie cellulaire, Hôpital Maisonneuve-Rosemont, Canada and Department of Medicine, Faculty of Medicine, Université de Montréal, Canada; 13 The Leucegene project at Institute for Research in Immunology and Cancer, Université de Montréal, Montréal, Canada; 14 Department of Biochemistry and Molecular Medicine, Faculty of Medicine, Université de Montréal, Montréal, Canada; 15 Queen Mary University of London, London, United Kingdom; 16 Barts Health NHS Trust, London, United Kingdom; 17 Biotech Research and Innovation Centre (BRIC), University of Copenhagen, Copenhagen, Denmark; 18 Department of Haemato-Oncology, St Bartholomew's Hospital, Barts Health NHS Trust, London, United Kingdom; 19 Experimental Hematology Lab, IIS-Fundación Jimenez Díaz, UAM, Madrid, Spain; 20 The Francis Crick Institute, London, United Kingdom; 21 Acute Leukaemia Team, Institute of Cancer Research, London, United Kingdom; 22 Haematopoietic Stem Cell Lab, The Francis Crick Institute, London, United Kingdom

**This supplemental material includes:**

- Supplemental tables 2, 9, 10
- Supplemental figures  1-4
- Supplemental Material & Methods

#### Supplemental Table 2

| Patient ID | Karyotype | Gene | Mutation | VAF |
|---|---|---|---|---|
| **P1** | 46,XY,t(6;9)(p23;q34) | *RUNX1* | c.C1192T:p.P398S | 12.92 |
| | | *STAG2* | c.A421G:p.N141D | 39.13 |
| | | *FLT3* | FLT3-ITD (72 bp) | 5.32 (allelic ratio) |
| **P2** | 46,XX,t(6;9)(p23;q34) | *FLT3* | FLT3-ITD (30 bp and 33 bp) | 0.8 and 1.16 (allelic ratio) |
| **P3** | 46,XY,t(6;9)(p23;q34) | *STAG2* | c.C412T:p.H138Y | 18.4 |
| | | *STAG2* | c.1027_1035TTAAGACAA | 13.2 |
| | | *FLT3* | FLT3-ITD (63 bp) | 0.32 (allelic ratio) |
| **P4** | 46,XY,t(6;9)(p23;q34) | *FLT3* | FLT3- ITD (54 bp and 111 bp) | 0.32 (allelic ratio) |

#### Supplemental Table 9

| Gene | Forward Primer | Reverse Primer | Amplicon length (bp) |
|---|---|---|---|
| *EYA3* | ATGCAGAGAGCACCACATTAG | TAGAAGGGTCTCCAGAGGAAAG | 105 |
| *SESN1* | CGAGATGAAGAAGAGGCAAGTC | GCTCTTGCTGGTGTAACTTCT | 113 |
| *PRDM2* | GACTCACCAGCATGGAGTTT | CAGAACTAGCTGTCCACTCTTTAG | 103 |
| *NFIX* | GCCACATCACATTGGAGTCA | CTCCTTGCTGGTTTGAACTATCT | 104 |
| *FOXC1* | ACCCGGACTCCTACAACAT | TCCTTCTCCTCCTTGTCCTT | 97 |
| *GGT5* | ATGCTGGTGGAGGACATTG | GGTGGTGAGTACAGGGTATAGT | 125 |
| *HOXA1* | CTTCTCCAGCGCAGACTTT | GTAGCCGTACTCTCCAACTTTC | 80 |
| *HOXA2* | GTTTCAGAACCGGAGGATGAA | TCCTCGTCCTCCTCTACTTTC | 114 |
| *HOXA3* | GCTGGAGAAAGAGTTCCACTTC | GTACTTCATGCGGCGATTCT | 124 |
| *HOXA4* | GGAGAAGGAGTTCCACTTCAAT | GGTCTTTCTTCCACTTCATCCT | 131 |
| *HOXA5* | TGCGCAAGCTGCACATA | GGTTGAAGTGGAACTCCTTCTC | 114 |
| *HOXA6* | TTGGATGCAGCGGATGAA | TAGCGGTTGAAGTGGAACTC | 123 |
| *HOXA7* | GAGGCCAATTTCCGCATCTA | CGGTTGAAGTGGAACTCCTT | 119 |
| *HOXA9* | GTGGTTCTCCTCCAGTTGATAG | AGTTGGCTGCTGGGTTATT | 123 |
| *HOXA10* | CCTTCCGAGAGCAGCAAA | GCTTCTTCCGACCACTCTTT | 103 |
| *HOXB2* | GCTGGAACTGGAGAAGGAAT | CCGGTTCTGAAACCAGACTT | 118 |
| *HOXB3* | GAAGTACAAGAAGGACCAGAAGG | CATGGAGTGTAAGGCGTTCAT | 124 |
| *HOXB4* | CAAAGAGCCCGTCGTCTAC | CGGTTGTAGTGAAATTCCTTCTC | 147 |
| *HOXB5* | GGGCAGACTCCGCAAATA | TAGCGGTTGAAGTGGAACTC | 143 |
| *HOXB6* | CGAGACAGAAGAGCAGAAGTG | CGTCTGGTAACGTGTGTATGT | 127 |
| *18S rRNA* | AAACGGCTACCACATCCAAG | CCTCCAATGGATCCTCGTTA | Endogenous control |
| *GAPDH* | CCATCACCATCTTCCAGGAG | GAGATGATGACCCTTTTGGC | Endogenous control |

### Supplemental Table 10

| Gene | Forward Primer | Reverse Primer | Amplicon length (bp) |
|---|---|---|---|
| ***FOXC1-1*** | TAATCGCTGCGTTCATCTCC | AGATTTAAGTGGTCTCCCTGAAC | 101 |
| ***FOXC1-2*** | GGCAGCGGAGCTGTTAAAT | GAATCCCAAGTCGTTCACTAGAAG | 92 |
| ***HOXA-1*** | TCTGGCGTCTAGGGACTG | CAGCTGGGTGGCCTTTAG | 98 |
| ***HOXA-2*** | GGGCAAGGCCTAGGAAA | CCCATACGGCTGTAATCAGT | 104 |
| ***HOXB-1*** | ACAGGCTCAGTGCCTTTGA | CATATTTATTTAATCACCCGGGAGTGG | 77 |
| ***HOXB-2*** | AGACCCAGGTCAGACTCTC | CCACAGTCTTCCACTCTACAAG | 90 |
| ***HOXB-3*** | ACAGCTCCAATCCTCTCTACTC | GGCTGCCTGCAGAAGTG | 90 |
| ***HOXB-4*** | CCATTGGCCTCAGAGAAACA | GAGGTGACAGCTGGAGATTAAG | 146 |
| **Positive control region** | AACTTCCTTGTGTTGTGTGTATTC | GAACTCAAAGCGGCTGAAATC | 117 |
| **ACTB (Negative control)** | CACCTGCACTCTGGGTAAG | TGAGGCCAAGTGTGACTTT | 127 |

#### Supplemental Figure 1

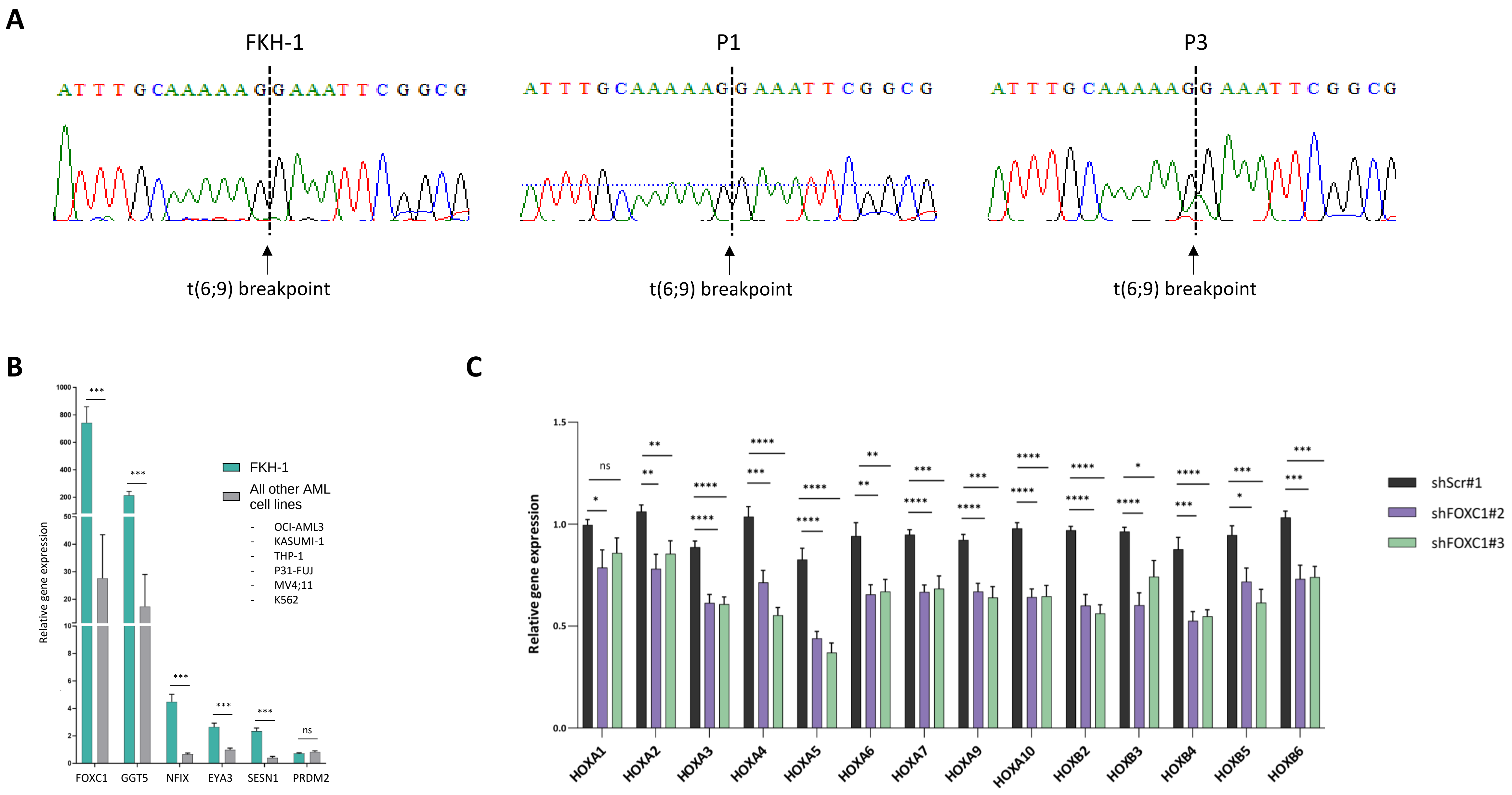

**Supplemental Figure 1**

**(A)** Sanger-sequencing chromatograms corresponding to the t(6;9)/DEK-NUP214 breakpoint region in the cDNA of two t(6;9) primary samples (P1 and P3) and the t(6;9) AML cell line FKH-1**.** Arrows indicate the exact point of the identical DEK::NUP214 breakpoint in the three samples. **(B)** Bar chart representing mRNA expression levels of *FOXC1*, *GGT5*, *NFIX*, *EYA3*, *SESN1*, and *PRDM2* assessed by RT-qPCR across the panel of AML cell lines: FKH-1, OCI-AML3, KASUMI-1, THP-1, P31-FUJ, MV4;11, and K562. Relative expression levels were calculated using the ΔCT method, normalized to the average of GAPDH and 18S rRNA. **(C)** Bar chart representing mRNA expression levels of *HOXA* and *HOXB* genes in *FOXC1* knockdown/scramble FKH-1 cells assessed by RT-qPCR. Relative expression levels were calculated using the ΔCT method, normalized to the average of GAPDH and 18S rRNA.

Statistical significance is indicated by asterisks (*: $p<0.05$, **: $p<0.01$, ***: $p<0.001$, ****: $p<0.0001$ and 'ns' denotes non-significant differences).

### Supplemental Figure 2

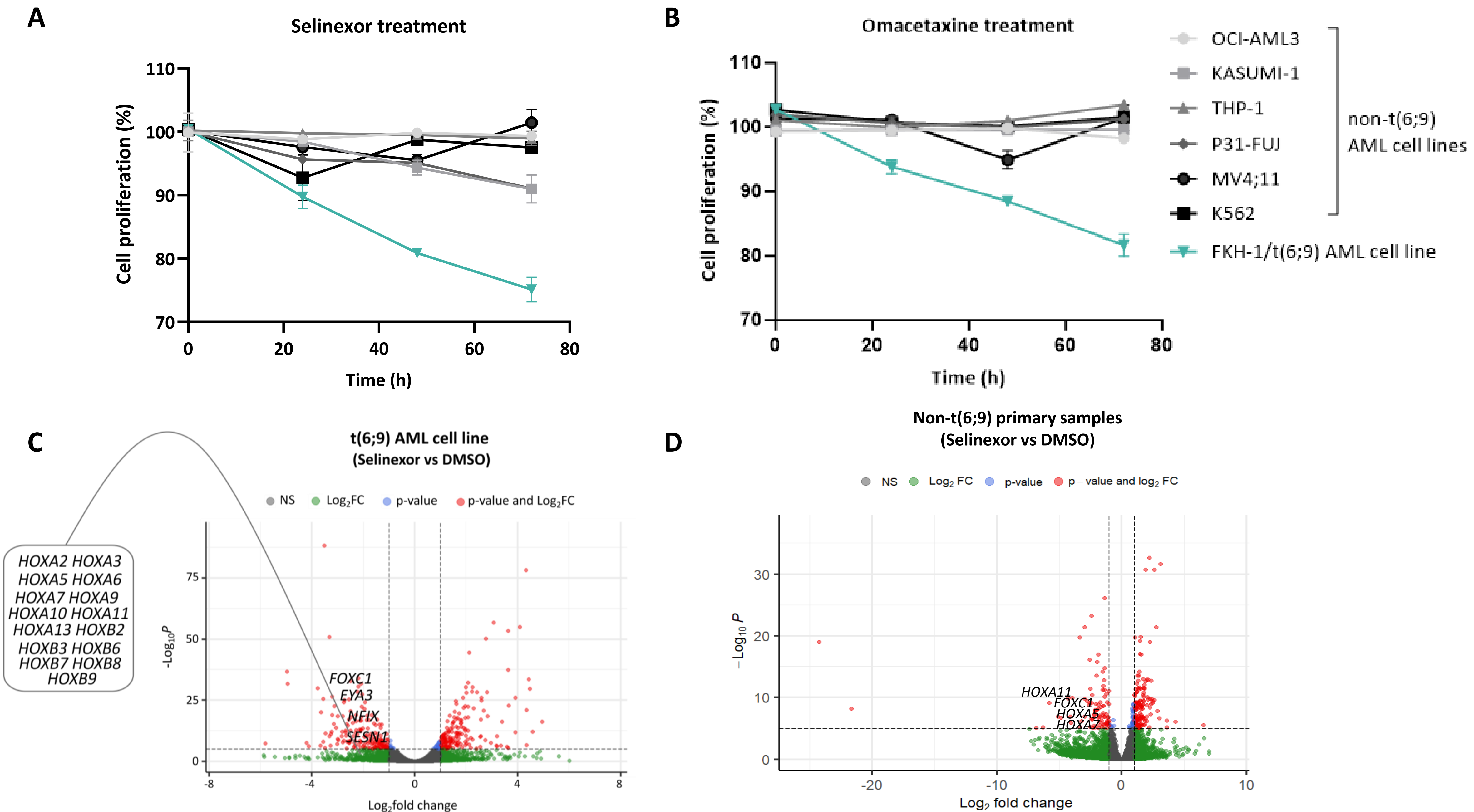

**Supplemental Figure 2.**

**(A) and (B)** Cell proliferation assays in a panel of AML cell lines treated with 200nM Selinexor (A) and 7nM Omacetaxine (B) for 72 hour. The panel includes the t(6;9) AML cell line FKH-1 (highlighted in green) and non-t(6;9) AML cell lines- OCI-AML3, KASUMI-1, THP-1, P31-FUJ, MV4;11, and K562- highlighted in shades of grey. **(C) and (D)** Volcano plots representing differentially expressed genes in t(6;9) AML cell line FKH-1 (C) and non-(6;9) primary samples (D) treated with 200nM Selinexor vs DMSO treatment control (RNA-seq analysis).

### Supplemental Figure 3

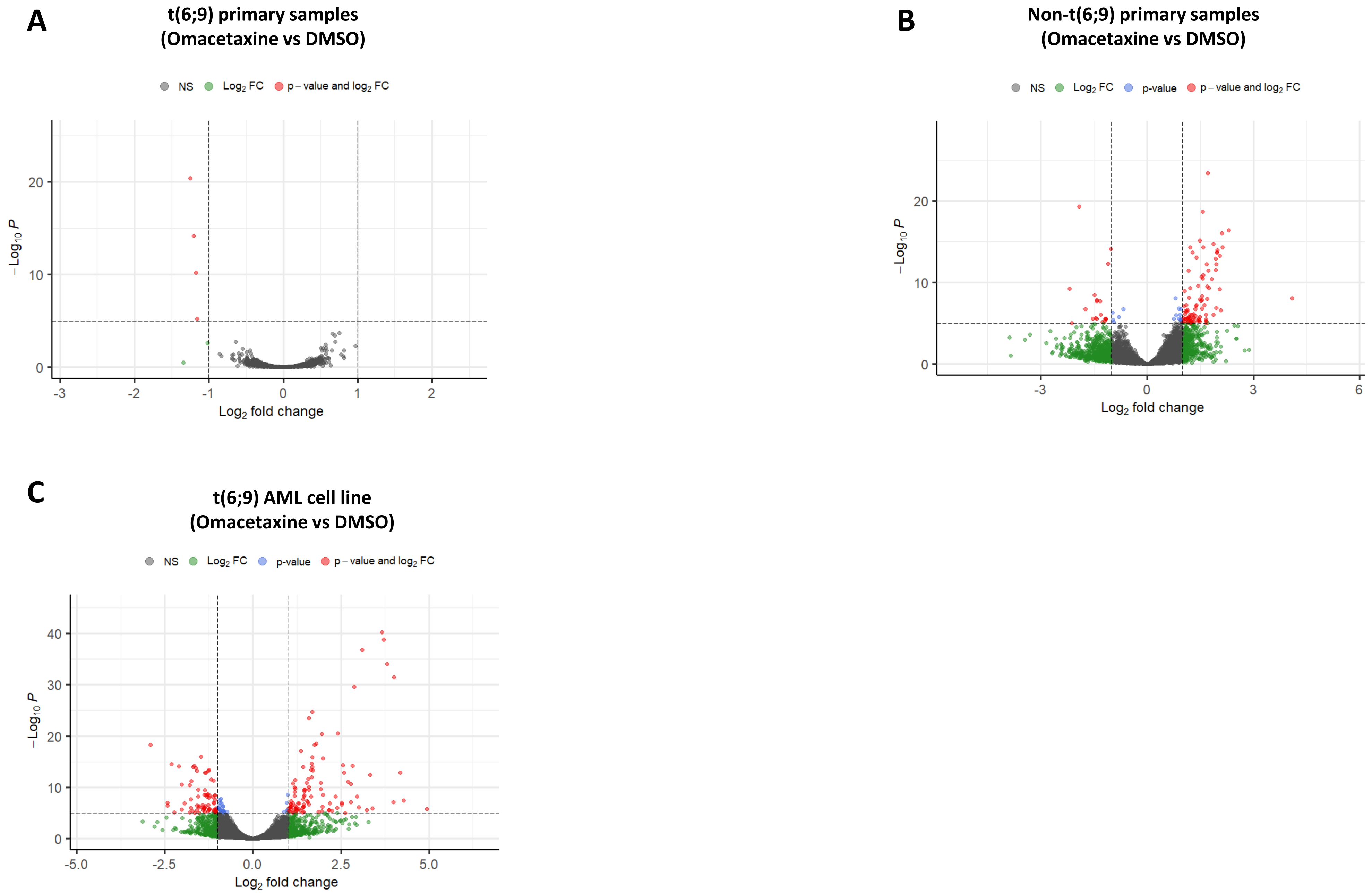

**Supplemental Figure 3.**

**(A)**, **(B) and (C)** Volcano plot representing differentially expressed genes in t(6;9) primary AML samples (A), non-(6;9) primary samples (B) and FKH1 cell line (C) treated with Omacetaxine (3 nM for primary samples and 7 nM for cell line) vs DMSO treatment control (RNA-seq analysis).

### Supplemental Figure 4

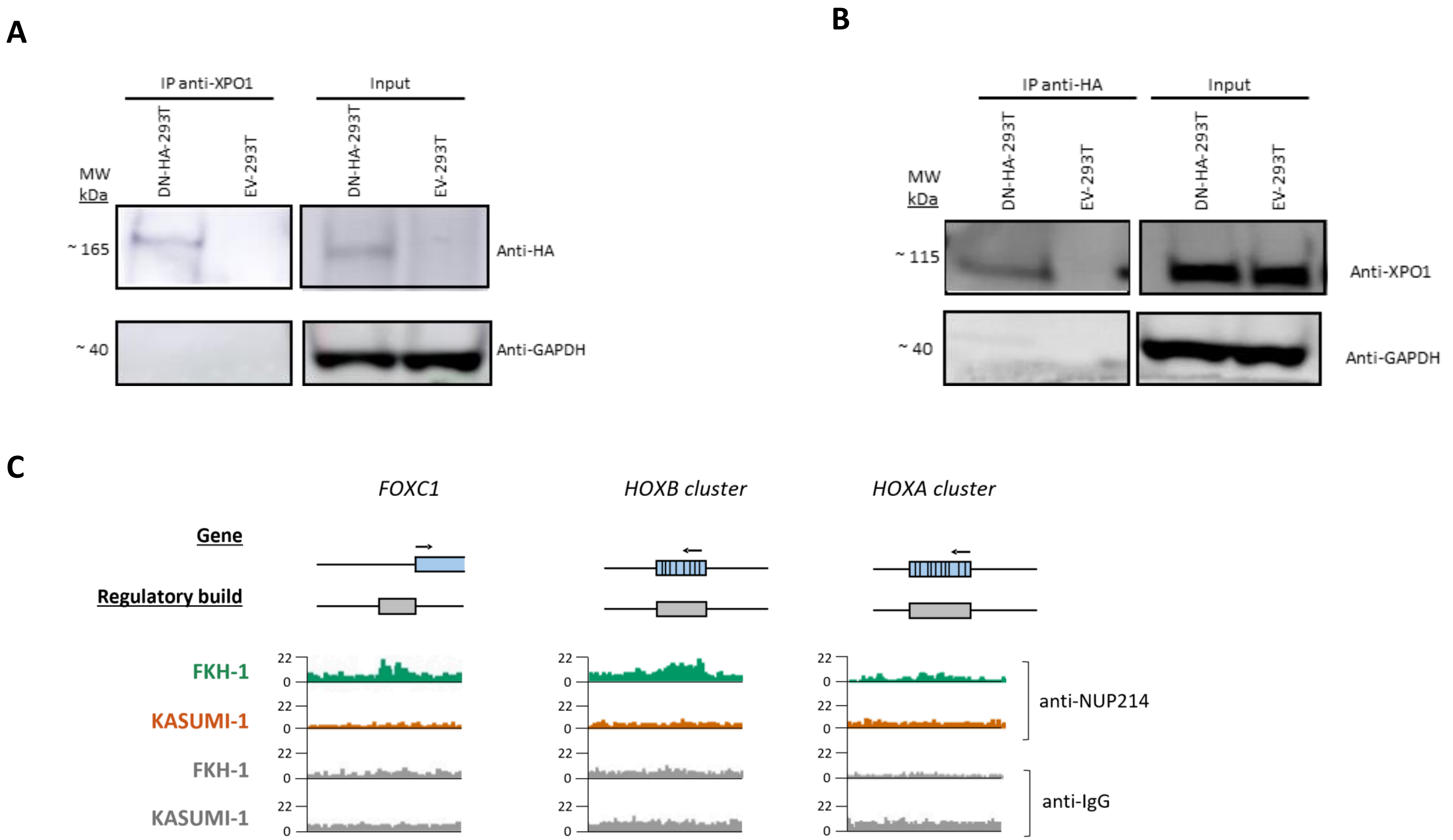

**Supplemental Figure 4.**

**(A) and (B)** Co-Immunoprecipitation (Co-IP) experiments in DN-HA-293T and EV-293T using anti-XPO1 (A) and anti-HA (B) antibodies. Input correspond to the unbound protein extracts. GAPDH was used as a loading control in western blotting. **(C)** IGV genome browser plots representing the binding peaks on the *FOXC1*, *HOXB* and *HOXA* promoters corresponding to the CUT&RUN experiments performed in FKH-1 and KASUMI-1 cells using NUP214 antibody (and anti-IgG as a control).

### Supplemental Material & Methods

#### Primary samples and cell lines culture

AML patients gave informed consent for the storage and use of their blood cells for research purposes. Experiments were performed in accordance with the Local Research Ethics Committee, as previously described [1]. Mononuclear cells from peripheral blood or bone marrow biopsies from the patients were isolated at the Barts Cancer Institute (QMUL) tissue bank facilities and stored in liquid nitrogen. Patients' clinical characteristics are listed in Supplementary Table 1. Primary human AML samples were thawed and co-cultured on MS5 stromal cells in Myelocult media (StemCell Technologies, H5100) with TPO, IL3, G-CSF (PeproTech, UK), and PenStrep (Gibco, UK).

AML cell lines FKH-1, OCI-AML3, and KASUMI-1 were cultured in RPMI medium supplemented with 20% FBS and 1% PenStrep (Gibco, UK). THP-1, P31-FUJ, MV4-11 and K562 were cultured in RPMI medium supplemented with 10% FBS and 1% PenStrep (Gibco, UK). HEK-293T cells were cultured in DMEM with 10% FBS and 1% PenStrep (Gibco, UK).

#### Reverse transcription (RT) and quantitative PCR

Total RNA was extracted from primary samples and cell lines using the RNeasy® Plus Mini Kit (Qiagen, Germany) and reverse-transcribed using a High-Capacity cDNA Reverse Transcription Kit (Applied Biosystems, Vilnius, Lithuania). Quantitative PCR (qPCR) was performed using cDNA and the Universal SYBR® Green (Bio-Rad, CA, USA) in the CFX384TM Real-Time System (Bio-Rad). All primers used are listed in Supplemental Table 9. All samples were run in triplicates and the ΔCT method was employed to normalize gene expression to the endogenous controls glyceraldehyde-3-phosphate dehydrogenase (GAPDH) and 18S ribosomal ribonucleic acid (18S rRNA).

#### Conventional PCR and Sanger sequencing

To amplify the DEK-NUP214 breakpoint, 1 µL of cDNA was mixed with 4 µL ReddyMix (2X, Thermo Fischer Scientific), 0.5 µL forward primer [GCATCTGTGAGGTTCTTGATTTG], 0.5 µL reverse primer [GGACAGTTTGGCTGGTACTT], and 4 µL nuclease-free water per 0.2 mL PCR tubes. The samples were subjected to the following thermal cycling conditions in a thermal cycler: initial denaturation at 95°C for 15 minutes, followed by 35 cycles of denaturation at 95°C for 30 seconds, annealing at 58°C for 30 seconds, and extension at 72°C for 30 seconds. A final extension at 72°C for 5 minutes was followed, and samples were held at 4°C until use. The PCR products were run on a 1.5% agarose gel at 100 V for 30 minutes. Bands of interest were excised and purified using a gel extraction kit (Qiagen), following the manufacturer's instructions.

The gel-extracted DNA was divided into two tubes and combined with 2.5 µL of corresponding forward and reverse primers. Samples were labelled with barcodes and sent to Eurofins (Germany) for Sanger sequencing. Sequencing results were visualized and manually reviewed using BioEdit software. The sequence was compared to the in silico-designed DEK-*NUP214* fusion sequence to pinpoint the breakpoint and validate the fusion junction.

#### Protein extraction and Western blotting

Cells were lysed with Pierce™ Radioimmunoprecipitation assay (RIPA) lysis buffer (ThermoFisher Scientific, USA) supplemented with protease inhibitor (Roche), phosphatase inhibitor (Roche), and benzonase® nuclease (Sigma). Protein quantifications were performed by BCA (Bicinchoninic acid) assay (Thermo Scientific, USA). Next, proteins from each cell line were resuspended in NuPAGE® LDS sample buffer 4X (Invitrogen, Carlsbad, CA) with NuPAGE® sample reducing agent 10X (Invitrogen, Carlsbad, CA) and heated at 95 °C for 5 minutes, prior being loaded on SDS-PAGE, NuPAGE™ 4-12% Bis-Tris (<150 kDa proteins) or 3-8% Tris-Acetate Gel (>150kDa proteins) (Invitrogen, CA), and ran in 1X 3-(N-morpholino)propanesulfonic acid (MOPS) or 1X Tris-Acetate Buffer (TAB) (Invitrogen) at 120 V for one hour. Proteins were transferred by dry transfer to nitrocellulose membrane (Invitrogen, Israel). After blocking with either 5% Bovine Serum Albumin (BSA) (Sigma) or 5% non-fat milk in TBST (1X Tris-Buffered Saline, 0.1% Tween 20 Detergent) for 1 hour at room temperature, the membranes were developed at 4 °C overnight with primary antibodies (anti-FOXC1 (1:1000 dilution, Abcam, ab227977); anti-XPO1/CRM1 (1:100 dilution, Santa Cruz Biotechnology, sc-74454); anti-HA (1:1000, Cell Signaling Technology, #2367); anti-GAPDH (14C10) (1:5000 dilution, Cell Signaling Technology, #2118)) in TBST containing either 5% BSA or 5% non-fat milk, according to manufacturer's instructions. The following day, membranes were washed three times in TBST and incubated with the corresponding secondary antibody for an hour (1:4000 dilution; Polyclonal goat anti-rabbit immunoglobulins/HRP P0448, or Polyclonal rabbit anti-mouse immunoglobulins/HRP P0260, DAKO). GAPDH was used as a loading control. Bands were revealed by incubation with ECL detection reagent (Cytiva Amersham, RPN2236) and developed chemiluminescence by Amersham Imager 600 (GE healthcare).

#### Drug treatments and cell viability assessment

Drug sensitivity and resistance-testing platform (DSRT) was used as previously performed in our t(6;9) patients [1-3]. We integrated our results with 141 AML samples treated with the same drug panel and spanning other cytogenetic groups (13 from our poor-risk cohort [1], 99 from Malani D. et al [2], 16 from Chen Y. et al. [3] and 17 unpublished samples listed in Supplementary table 4).

Cell viability assays were performed over 72 hours at increasing drug concentrations of Selinexor (1, 10, 100, 500, 1000, and 1500 nM) and Omacetaxine (1.5625, 3.125, 6.25, 12.5, 25, and 50 nM) in the panel of AML cell lines (FKH-1, OCI-AML3, KASUMI-1, THP-1, P31-FUJ, MV4-11, and K562). 40,000 cells from FKH-1, 15,000 cells from the other AML cell lines (OCI-AML3, KASUMI-1, THP-1, P31-FUJ, MV4-11, K562), and 10,000 primary patient cells were seeded per well into three technical replicates in a 96-well plate. Cell viability was assessed by CellTiter-Glo® (CTG) (Promega) assay according to the manufacturer's instructions, which allows quantitative measurement of ATP to determine cell viability. Next, luminescence readings were obtained to assess cell viability. The luminescence values were first normalized to the control wells to account for background signals. The normalized viability data for each drug concentration were then imported into GraphPad Prism software. Dose-response curves were generated, and IC50 values were determined by fitting the data to a non-linear regression model (log[inhibitor] vs. normalized response). The IC50 represents the concentration at which 50% inhibition of cell viability was observed for each drug in the panel of AML cell lines.

Drug sensitivity tests were conducted utilizing the Guava® ViaCount™ Flex kit (SKU 4500-0110), which distinguishes between viable and nonviable cells based on their permeability to specific DNA-binding dyes, enabling precise quantification of live and dead nucleated cells in suspension. Cell viability was evaluated over 72 hours with 200nM Selinexor, 7nM Omacetaxine, or DMSO across a panel of AML cell lines (FKH-1, OCI-AML3, KASUMI-1, THP-1, P31-FUJ, MV4-11, and K562). Cells were plated at 40,000 cells per well in triplicate in a 96-well plate. The Guava® ViaCount™ Flex assay was employed to assess cell viability at 0h, 24h, 48h, and 72h, according to the manufacturer's protocol, using the Guava EasyCyte HT flow cytometer, with live cells manually gated and measurements taken in triplicate.

#### RNA sequencing

Primary patient samples from two t(6;9) patients (patients P1 and P3) and two non-t(6;9) AML patients (patients P13 and P28), and FKH-1 cells were seeded (300,000 cells / well) into two technical replicates in 6-well plates pre-seeded with irradiated MS5 stroma cells, and treated for 48 hours with 200 nM Selinexor, Omacetaxine (3 nM for primary samples and 7 nM for cell lines), and DMSO as a control , followed by RNA extraction. RNA concentrations were determined using Qubit® 3.0 Fluorometer. All samples were checked for RNA integrity and quality control and sequenced by Novogene. Poly A enriched mRNA library was constructed and sequenced by Illumina PE150 to generate 20 M reads, 6 G of raw data per sample. Raw paired FASTQ reads were aligned to GRCh38.91 using HiSat2 (version 2.1.1). Feature counting was carried out at the gene level using htseq-count (version 0.13.5). Genes were filtered prior to downstream analysis with only genes that had more than 10 reads in at least four samples being retained (16,435 genes). Differential gene expression analysis was carried out for each drug separately using DESeq2 (version 1.42), comparing the drug samples to the DMSO samples. For samples with paired replicates the replicated was added to the design matrix (~ replicate + treatment). Gene set enrichment analysis (GSEA) was run using the fgsea R package (version 1.28.0).

Leucegene RNAseq libraries were constructed according to TruSeq Protocols (Illumina) and sequencing was performed using either an Illumina HiSeq 2000/4000 or a NovaSeq 6000 instruments (Geo SuperSeries GSE232130). Mapping was performed using STAR v2.7.1 and genes expression levels were quantified using RSEM v1.3.2 (PMID 21816040). Leucegene cohort and our poor-risk AML cohort data were batched-corrected on the read counts using the Lima-voom R package (PMID 25605792). EPCY was used to identify differentially expressed transcripts that are predictive of t(6;9) status (https://github.com/iric-soft/epcy).

#### *FOXC1* silencing

Two lentiviral shRNA vectors EGFP-tagged targeting *FOXC1* (shFOXC1#2 [5'-ACTCTCCAGTGAACGGGAATA-3'] and shFOXC1#3 [5'-GAGCTTTCGTCTACGACTGTA-3']) and non-target Scramble control (shScr#1 [3'-CCTAAGGTTAAGTCGCCCTCG-5']) were purchased from VectorBuilder (Chicago, USA). Viral particles for all shRNAs were produced by transient $CaCl_2$ transfection of HEK-293T cells. At 48-hour post-transfection, viral particles were harvested and concentrated by ultracentrifugation at 22,000 rpm at 4°C for 2 hours. The supernatant was removed and resuspended in 300 µL of fresh media, then agitated at 4°C for 2-4 hours. For viral particle quantification, 50,000 HEK-293T cells per well were seeded in 500 µL of DMEM supplemented with 10% heat-activated FBS and 1% PenStrep (Gibco, UK) in a 24-well plate. Gradually increasing volumes of the agitated viral media (1:200) (2, 4, 8, 16, 32, and 64 µL) were added to the wells. After four days, viral titters were assessed using flow cytometry based on EGFP expression to determine the proportion of transduced cells.

Viral particles at a multiplicity of infection (MOI) of 10 were added to the FKH-1 cells. The cells were then incubated overnight at 37 °C with 5% $CO_2$. The following day, cells were washed and resuspended in fresh media 4 days after transduction, cells were sorted based on EGFP expression (FACS Aria Fusion Cytometer Analyser, BD, Biosciences, CA, USA).

#### Colony forming unit (CFU) assay

shRNA-transduced (10,000 cells per plate) were seeded into triplicates immediately after EGFP cell sorting in 1.2 mL MethoCult Classic (#H4434, StemCell Technologies, Canada) supplemented with 1% PenStrep (Gibco, UK) in 35 mm x 10 mm dishes and incubated at 37 °C under hypoxic condition (3% $O_2$) for 14 days. At the end of this period, colonies were scored and analysed under the microscope.

#### Cell cycle and apoptosis assays

Three days after EGFP cell sorting, shRNA-transduced cells were harvested and washed. For cell cycle assay, cells were fixed in 250 μL of Fixation/Permeabilisation solution (Cytofix/Cytoperm™, #554714, BD Biosciences) and incubated at 4 °C for 20 minutes. Next, fixed cells were washed twice in 1X BD Perm/Wash™ buffer (Cytofix/Cytoperm™, #554714, BD Biosciences). Following the washing step, cells were resuspended in BD Perm/Wash™ buffer with DAPI (1/100) and incubated at 4°C for 30 minutes in the dark and analysed based on DNA content by FACS (BD LSRFortessa, USA).

For apoptosis assay, cells were incubated with Annexin V solution (1X Annexin V buffer with Annexin V antibody (Alexa Fluor® 647Annexin V, Cat:640912, Biolegend) at room temperature in the dark for 15 minutes. Beads for compensation were prepared by incubating 1 drop of beads (OneComp eBeads, Invitrogen, Carlsbad, CA) in DPBS (Gibco, UK) with APC antibody (APC anti-human CD11b, Clone: ICRF44, Biolegend) in the dark for 15 minutes. Then, samples were washed with FACS buffer (DPBS supplemented with 2% FBS and 1% PenStrep) and resuspended in FACS buffer with DAPI prior analysis by FACS (BD LSRFortessa, USA).

#### DEK::NUP214 constitutive model

To generate human HEK-293T cells with stable overexpression of the DEK-NUP214 fusion protein, a lentiviral vector encoding DEK::NUP214 (pLV[Exp]-EGFP-hPGK>{HA-DEKNUP214-FLAG}) (DN-HA) along with an empty vector control (pLV[Exp]-EGFP-hPGK>ORF_Stuffer) (EV) were ordered from VectorBuilder (Chicago, USA). Lentiviral particles were produced as described before and HEK-293T cells were transduced to establish stable expression of the fusion protein, as proven by Western blotting with anti-HA antibody. HEK-293T cells expressing DEK::NUP214 (DN-HA-293T) were utilized for subsequent experimental analyses using HEK-293T transduced with the empty vector (EV-293T) as a control.

#### Co-Immunoprecipitation (Co-IP)

DN-HA-293T cells were washed in DPBS twice and then resuspended cell pellet in ice-cold Lysis Buffer (50 mL of 1M HEPES pH 7.0-7.6, 3.75 mL of 4M NaCl, 200 µL of 0.5M EDTA, 100 µL of NP-40, $dH_2O$ up to 100 mL) at 50 µL per million cells. Lysed cells were incubated at 4 °C for 15 minutes and then centrifuged at 15,000 x g for 10 minutes at 4°C. The supernatant was removed and transferred to a new tube. 50 µL of Pierce™ Protein A/G magnetic beads (88802, ThermoFisher) were added per sample to an Eppendorf tube and washed with 1 mL of ice-cold Lysis-PI buffer (10 mL of Lysis Buffer, 1 EDTA-free Protease inhibitor cocktail tablet (A32955 ThermoFisher)) three times. Beads then were resuspended in 30 µL ice-cold Lysis-PI buffer and added to cell lysate following 15 minutes incubation in an orbital shaker at 4°C.  Beads were collected with a magnetic stand and then removed the lysate flow-through, and the supernatant was transferred to a new pre-cooled centrifuge tube. After protein quantification, 500 µg of protein was taken and added 25 µL of anti-XPO1 (Santa Cruz Biotechnology, sc-74457) or 30 µL of anti-HA (Cell Signaling, #2367) per condition and the volume was made up to 500 µL total using Lysis Buffer. IP reaction was incubated overnight at 4°C with mixing. Following day, 50 µL of Pierce Protein A/G magnetic beads were placed into a 1.5 mL microcentrifuge tube for each IP reaction. 175 µL of Wash Buffer (10 mL 10X TBS, 90 mL H2O, 50 µL Tween-20, dH20 up to 100 mL) to the beads and gently vortexed to mix. The tube was placed into a magnetic stand to collect the beads. After two washes, the IP mixture was added to the beads and incubated for 1 hour at 4°C with mixing. The beads were collected and washed twice with Wash Buffer and purified water. 55 µL of Low-pH Elution Buffer (750.7 mg glycine, 100 mL of dH2O made to pH 2.0 using HCl) was added to the tube and incubated at room temperature for 10 minutes with mixing. The beads were magnetically separated and the supernatant containing target antigen was saved and 8.25 µL of Neutralisation Buffer (1 M Tris pH 7.5-9) was added to the each elute to neutralise the low pH. Western blotting was performed as described above on immunoprecipitated samples (DN-HA-293T and EV-293T for IP anti-XPO1 and anti-HA) and input using the following antibodies: anti-XPO1 (1:100 dilution, Santa Cruz Biotechnology, sc-74457), anti-HA (1:1000 dilution, Cell Signaling, #2367), or anti-GAPDH (1:5000 dilution, as a loading control).

#### CUT&Run assay

*Cleavage under targets and release using nuclease* Assay (CUT & Run) was performed according to the EpiCypher CUTANA CUT&Run protocol in DN-HA-293T and EV-293FT cells and FKH-1 and KASUMI-1 cell lines as previously described [4].

In brief, 500,000 cells were harvested per condition, and after two washes, cells were incubated with activated Concanavalin A (ConA) beads (Bangs Lab, BP531) for 10 minutes at room temperature. After removing supernatant, 0.5 µL corresponding antibody (anti-NUP214, ab70497 Abcam; anti-HA, 2367 Cell Signaling; or anti-IgG, ab171870 Abcam) was added and incubated overnight. The following day, samples were washed twice in cold Digitonin Buffer and incubated in 50 µL Digitonin Buffer with pAG-MNase for 10 minutes. Supernatant was removed and 50 µL Digitonin and 1 µL 100 mM $CaCl_2$ were added to the samples and incubated for 2 hours to activate and cleave targeted chromatin. Next, 33 µL of Stop Buffer were added to samples that were incubated at 37 °C for 10 minutes to release chromatin to the supernatant and to degrade the RNA. Supernatants were collected and purified by using CUTANA DNA purification kit (Zymo research, D4014) according to the manufacturer's instructions. Eluted DNA was quantified using Qubit™ fluorometer and used for library preparation using NEB Library Prep Kit (E7645). PCR enrichment of adaptor-ligated DNA was performed using NEBNext® Oligo Kit according to the manufacturer's instructions (NEBNext® Multiplex Oligos for Illumina® (Index Primers Set 1, 2, and 3)). After cleanup of amplified adaptor-ligated DNA, its integrity and concentration were checked using TapeStation (Agilent High Sensitivity D1000 Screen Tape System) according to the manufacturer's instructions. Following quality and concentration check, the libraries were sequenced on Novaseq 6000 platform using 150bp paired-end reads. CUT&RUN paired end reads were trimmed using Trim Galore and aligned to the hg38 genome assembly using Bowtie2 (v2.4.5). The obtained bam files were sorted, indexed, and used for generating bigwig files. Bigwig files were converted to bedGraph format using bigWigToBedGraph (version 448) and peaks were called using macs2 (version 2.2.7.1) bdgpeakcall. Peaks were annotated with the ChIPseeker R package (version 1.40.0). Binding peaks located in promoters (≤1 kb or 1–2 kb from the transcription start site) and with a peak score of ≥150 for the DN-293T and ≥110 in FKH-1 cells, while absent in EV-293T and Kasumi-1, respectively, were selected.
CUT&RUN sequencing was also performed after treatment with Selinexor for 48 hours at a concentration of 200nM in the FKH-1 and KASUMI-1 cells using anti-NUP214 (ab70497 Abcam).

CUT&RUN-qPCR was then used to further validate the CUT&RUN-sequencing results, following the experimental procedure outlined in a published STAR protocol [5]. We first performed CUT&RUN in the FKH-1 and KASUMI-1 cell lines as described earlier, and the eluted DNA was then used for quantitative PCR (qPCR) analysis to validate the DEK-NUP214 binding sites on the *FOXC1*, *HOXA*, and *HOXB* genes clusters. Primer pairs were designed for the identified target regions: *FOXC1* promoter (two pairs), *HOXA* cluster (two pairs), and the *HOXB* cluster (four pairs, as the identified binding region was longer). Additionally, a positive control was included, located at a region on chromosome 1, which showed consistent strong enrichment across all samples regardless of DEK-NUP214 presence. A negative control, located upstream of the beta-actin (*ACTB*) gene and showing consistent lack of binding across all samples, was also included to confirm the absence of binding. All primers used are listed in Supplementary Table 10.
Then, qPCR was performed using 0.5 ng/µL eluted DNA per condition and the Universal SYBR® Green (Bio-Rad, CA, USA) reagent in a CFX384TM Real-Time System (Bio-Rad). Each sample was run in triplicates, and the ΔCT method ($2^{-\Delta\Delta Ct}$ method) was applied to normalize binding enrichment to the positive and negative controls. Fold change was calculated relative to the FKH-1_IgG sample. Data from different primer pairs targeting the same promoter regions of a given gene were integrated to ensure robustness and accuracy of the data.

#### Statistics

Data were presented as mean of three biological replicates ± SEM using GraphPad v.8 (GraphPad Prism 8 Software) and analysed using unpaired Student's t-test p-values < 0.05 were considered to be statistically significant.

#### Data availability

The sequencing data generated in this study have been deposited in the Gene Expression Omnibus (GEO) database under accession code GSE279056.
